# Whole exome sequencing reveals a mutation in *ARMC9* as a cause of mental retardation, ptosis and polydactyly

**DOI:** 10.1101/109124

**Authors:** Anjana Kar, Shubha R Phadke, Aneek Das Bhowmik, Ashwin Dalal

## Abstract

Intellectual disability (ID) refers to deficits in mental abilities, social behaviour and motor skills to perform activities of daily living as compared to peers. Numerous genetic and environmental factors may be responsible for ID. We report on identification of a novel gene for syndromic ID, using homozygosity mapping followed by exome sequencing in a family with mental retardation, ptosis and polydactyly. The analysis revealed a synonymous mutation c.879G>A which leads to a splicing defect in *ARMC9* gene. The variant is present in conserved region of ARM domain of ARMC9 protein which is predicted to form a platform for protein interaction. This domain is likely to be altered in patients due to splicing defect caused by this synonymous mutation. Our study was helpful in elucidation of molecular basis of mental retardation, ptosis and polydactyly phenotype and addition of *ARMC9* to group of genes leading to syndromic ID.

## INTRODUCTION

Intellectual disability has a prevalence of 1-3% in population and can result from heterogeneous causes like environmental/nutritional effect, chromosomal or monogenic causes^1^. More than 230 genes have been reported to be involved in causation of syndromes of intellectual disability. Mental retardation, ptosis and polydactyly^2^ are a distinctive combination of clinical features reported by us. This is a rare type of intellectual disability syndrome, reported only in a single consanguineous Muslim family from India, where three individuals were affected^2^.

Intellectual disability, short stature and polydactyly was suggestive of possibility of Bardet Biedl syndrome (BBS, MIM: 209900), but absence of obesity, renal abnormality, retinopathy and normal sexual development were some of the phenotypes that were different from BBS. Another possibility was 3MC syndrome, which was formally known as Carnevale syndrome (MIM: 265050), however patients did not had hip dysplasia, cryptorchidism and abdominal muscle defect. For identification of candidate gene in this family we have employed homozygosity mapping followed by exome sequencing in all the three affected siblings with mental retardation, ptosis and polydactyly phenotype.

## MATERIALS AND METHODS

### Patients Details

We described the clinical features in three affected siblings born out of consanguineous union, characterized as mental retardation, ptosis and polydactyly phenotype^2^. The proband showed severe intellectual disability (ID), bilateral ptosis, downslanting palpebral fissures, hypertelorism, round face, high arched palate, clinodactyly, tapering of fingers and hypermobile patellae. He also had joint laxity in the metacarpophalangeal and interphalangeal joints in hands and short stature. Female sibling was also proportionately short with stocky build. She had moderate ID, ptosis in right eye, pointed chin, bilateral post axial polydactyly with joint laxity. The third sibling had marked joint laxity similar to proband, moderate ID, bilateral ptosis and bilateral post axial polydactyly (Table I). We collected blood samples from all affected siblings and both parents after informed consent. This study was approved by the Institutional Ethics Committee.

**Table I:** Clinical Features of patients with Mental retardation, ptosis and polydactyly syndrome.

| Phenotype | II- 1 | II-2 | II-3 |
| --- | --- | --- | --- |
| Intellectual disability | severe | Moderate | Severe |
| Ptosis | Bilateral | Right Eye | Bilateral |
| Polydactyly (Hand) | - | + | - |
| Polydactyly (Feet) | - | + | + |
| Clinodactyly | + | - | - |
| Downslanting palpebral fissure | + | + | + |
| Short Stature | + | + | + |
| Stocky build | NA | + | NA |
| Pointed chin | - | + | - |
| Hypertelorism | + | NA | - |
| Hypermobile patellae | + | NA | NA |
| Joint laxity in metacarpophalangeal and interphalangeal joint | + | + | + |
| Round face | + | - | + |
| Tapering fingers | + | NA | -NA |
| Flat Feet | - | + | - |

### Genetic Testing

Genomic DNA from all affected and unaffected family members was extracted from whole blood. SNP array analysis in all three affected individuals was performed by using Illumina Human-CytoSNP-12 v.2.1 DNA Analysis BeadChip Kit (Illumina Inc., San Diego, CA). Analysis of array data was performed using Genome studio software. Size threshold for analysis was kept as 200 kb for deletion, 500 kb for duplication and 1 Mb for homozygous regions. Genomic Oligoarray and SNP evaluation tool (v 3.0) (http://firefly.ccs.miami.edu/cgi-bin/ROH/ROH_analysis_tool.cgi) was used for detection of known disease causing OMIM genes in the common homozygous regions.

Exome sequencing was performed in all three affected sibs. Agilent SureSelectXT V5 exome capture kit (Agilent Technologies, Santa Clara, CA) was used for preparation of exome library of using ~3 µg of genomic DNA. Exome library was sequenced to mean 100X coverage on Illumina HiSeq2000 sequencing platform. Alignment to human reference genome (GRCh37/hg19) and variant calling were performed using NEXTGENe v.2.3.4.4 and Annovar^3^, respectively. Variants with minor allele frequency of >0.01 in 1000 Genome project, EVS (Exome Variant Server), ExAC databases and in-house database were excluded in the study. Only nonsense, non-synonymous, splice site variants, synonymous splice site variants, insertion and deletion variants that affect coding regions of gene were used for interpretation. PCR was performed using gene specific primer set (listed in Supplementary Table SI) followed by Sanger sequencing on ABI3130 Genetic analyzer (Life Technologies, Carlsbad, CA) following the manufacturer’s protocol in control, parents, unaffected sib and affected sib to validate variants identified on exome sequencing.

### In vivo analysis of mutation at splice site

An observed variation was evaluated for splicing defect using pCAS2^4^ minigene construct. Wild type (ARMC9_Wt) and mutant (ARMC9_Mt) constructs were created by PCR amplification from control and patient sample respectively followed by cloning them in *BamH* I and *Mlu* I restriction site of pCAS2 vector. Recombinant constructs as well as pCAS2 (as control) were transfected into COS7 cells using Lipofectamine 2000 (Invitrogen, Carlsbad, CA) according to the manufacturer’s instructions. Cells were harvested after 18 hr of transfection and total RNA was extracted from each transfectant using RNeasy Mini Kit (Qiagen, Hilden, Germany), as per the manufacturer’s instruction. Total RNA was subjected to RT PCR using SuperScript III (Invitrogen, Carlsbad, CA) followed by plasmid specific amplification (P1 and P2 primer) and evaluated on 2 % agarose gel. Appropriate bands were purified using QIAquick Gel Extraction Kit (Qiagen, Hilden, Germany) and subjected to Sanger sequencing using on ABI3130 Genetic analyzer (Life Technologies, Carlsbad, CA).

## RESULTS

### Homozygosity mapping and Exome sequencing

We did not identify any significant copy number variations in any of the affected siblings. Regions of loss of heterozygosity (LOH) greater than 1 MB shared by affected siblings were identified on chromosome 2, 7, 10 and 14 (Supplementary Table SII), harbouring 228 Refseq genes and including 81 OMIM genes. The largest region of shared homozygosity was on chromosome 2q of 11 Mb, harbouring 141 genes. None of the genes in the shared regions were found to be associated with intellectual disability, ptosis and polydactyly occurring together. Whole exome sequencing was performed for all three affected siblings. Mapping and alignment to human reference genome was performed using NEXTGENe v.2.3.4.4 software (SoftGenetics, LLC, PA). Variants were filtered for minor allele frequency (MAF) ≤ 0.01 in 1000 Genome project, EVS, ExAC and in-house database of 30 Indian exomes.

Mutation databases such as HGMD and Clinvar were also by splice site prediction tools such as Human Splice Finder5 checked for presence of reported pathogenic variants in affected individuals. After filtering of variants, two candidate variants (c.654A>C in *UGT1A7* and c.879G>A in *ARMC9*) and MutPred Splice6. Sanger sequencing confirmed presence of homozygous c.879G>A mutation in all three affected siblings (Figure 1 B). The variant was found to be heterozygous present in shared homozygous region of 11 Mb on in both parents and absent in unaffected sibling. Chromosome 2 were studied further (Supplementary Table SIII and Table II).

**Figure 1:**
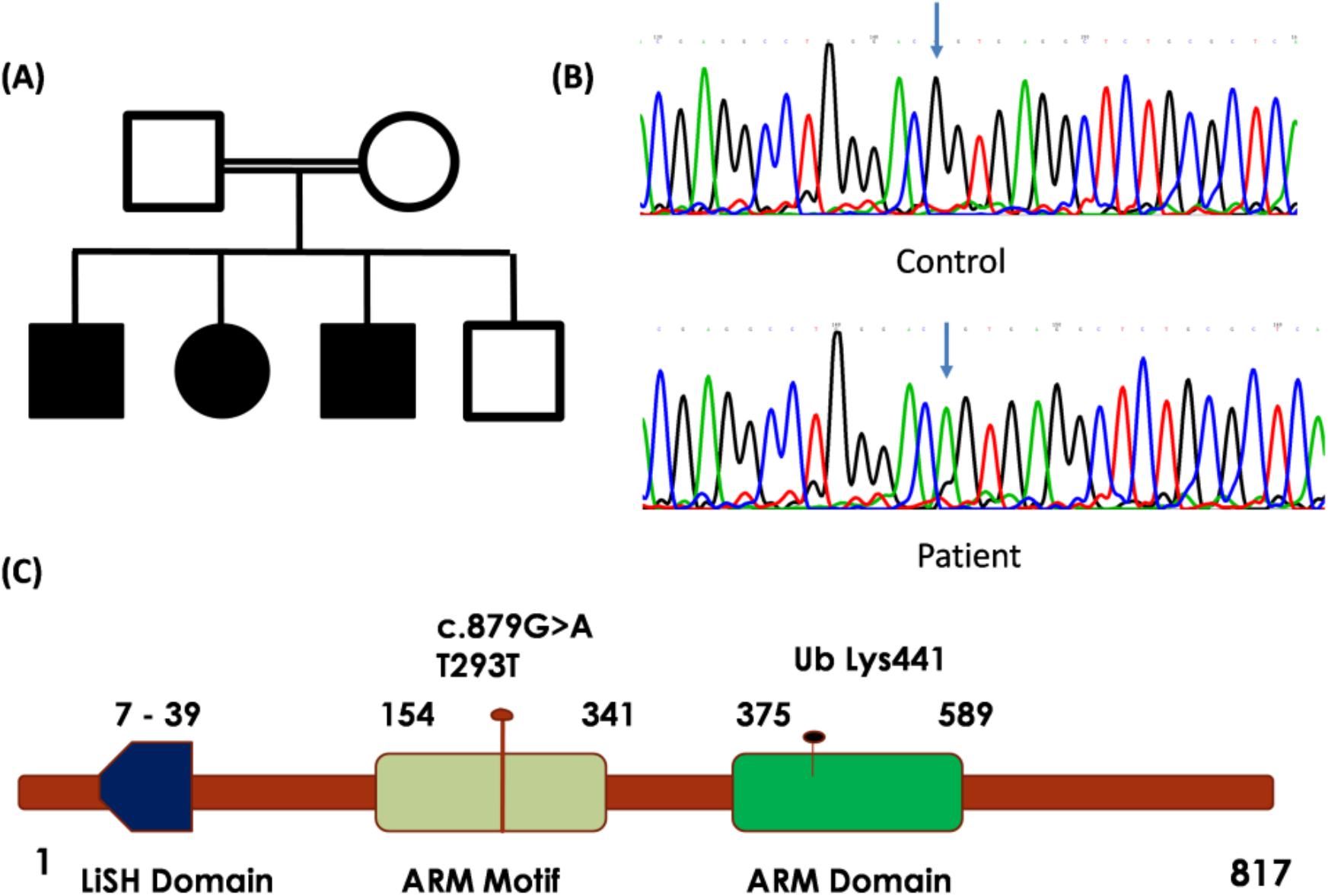
Mutation in ARM domain of *ARMC9* gene in patients with mental retardation, ptosis and polydactyly syndrome. (A) Pedigree of consanguineous family with three affected offspring affected with mental retardation, ptosis and polydactyly phenotype. (B) Sanger sequencing chromatogram of Control and patient showing c.879G>A indicated by arrows. (C) Schematic illustration of ARMC9 with location of mutation c.879G>A.

**Table II:** List of common variants in three affected patients II-1, II-2 and III-3

| S.No | Gene | Position | Variant | Protein variant | Gene Function |
| --- | --- | --- | --- | --- | --- |
| 1 | ARMC9 | Chr2:<br>232104754 | c.879G>A | p.Thr293Thr | Armadillo repeat containing 9<br>May be involved in microtubule dynamics |
| 2 | UGT1A7 | Chr2:<br>234591237 | c.654A>C | p.Leu218Phe | UDP Glucuronosyltransferase family 1<br>member A7<br>Enzyme of Glucuronidation pathway |
| 3 | ZFPM1 | Chr16:<br>88599697 | c.1331_1339delinsCCC | p.Phe446_449Leu | Zinc finger protein Multitype 1<br>Partner of hematopoietic transcription<br>factor GATA |
| 4 | SIGLEC12 | Chr19:<br>52004791 | c.193delinsAG | p.Ala66GlyFs*50 | Sialic acid binding immunoglobulin like<br>lectin 12<br>Cell surface protein of Ig superfamily |
| 5 | LILRB2 | Chr19:<br>54783375 | c.483A>C | p.Glu161Asp | Leukocyte immunoglobulin like receptor<br>subfamily B member 2 |

The variant in UGT1A7 was predicted to be pathogenic but the function of UGT1A7 gene in glucuronidation pathway did not appear to be related with the phenotype in the affected siblings. The variant *ARMC9* c.879G>A, on chromosome minigene. PCR amplification of pCAS2 minigene specific 2q37.1, was found to be homozygous in all affected siblings, present in shared homozygous region, absent in 1000 Genome database, EVS, ExAC, in-house database of 50 exomes and predicted to be damaging by various pathogenicity prediction software. This variant is a synonymous variant in the last base of exon 9 of *ARMC9* gene and was predicted to affect splicing by splice site prediction tools such as Human Splice Finder^5^ and MutPred Splice^6^. Sanger sequencing confirmed presence of homozygous c.879G>A mutation in all three affected siblings (Figure 1 B). The variant was found to be heterozygous in both parents and absent in unaffected sibling.

### Characterization of mutation by Minigene assay for splicing defect

Since *ARMC9* is not expressed in blood, in vivo characterization of this mutation was done to determine whether it causes a defect in mRNA splicing using pCAS2 minigene. PCR amplification of pCAS2 minigene specific primer (P1 and P2) from cDNA prepared using all transfectants were checked on 2% agarose gel, which revealed presence of 255 bp band in PCAS (control plasmid) and ARMC9_Mt whereas ARMC9_Wt had 354 bp fragment (Figure 2).

**Figure 2:**
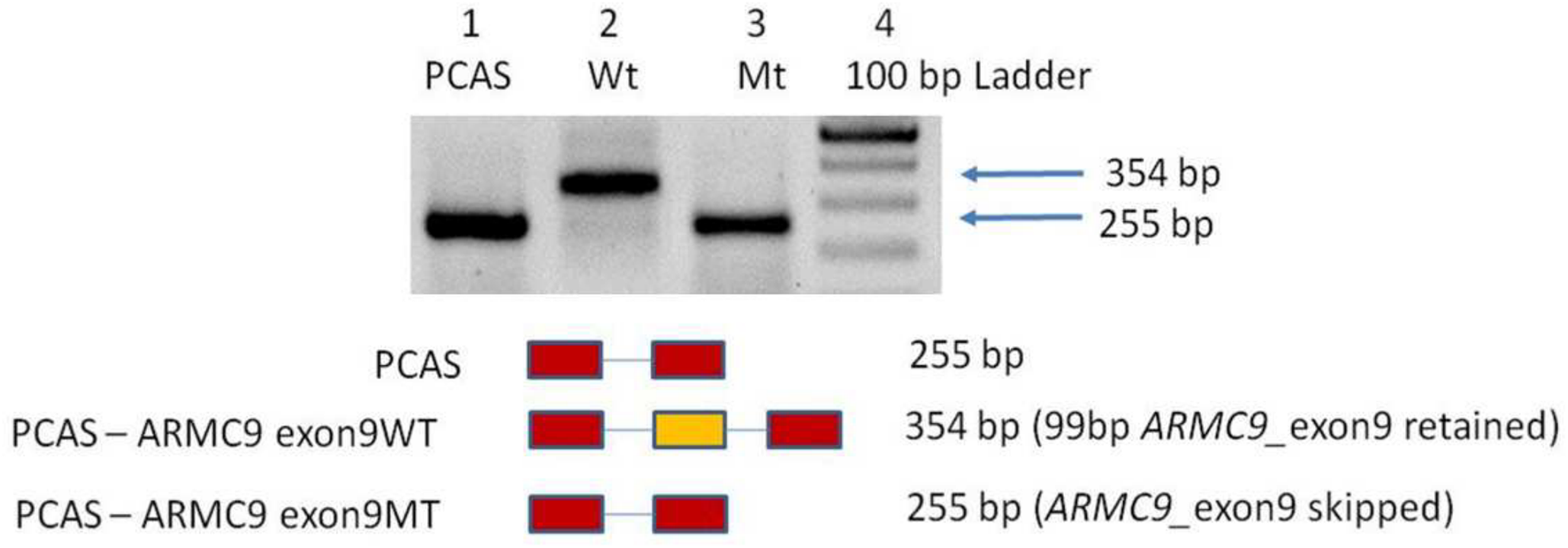
Molecular characterization of c.879G>A mutation. Results of RT-PCR performed on RNA isolated from transfected COS7 cells. Lane 1: empty pCAS2 Vector, lane 2 pCAS_ARMC9_WT vector, lane 3: pCAS_ARMC9_MT vector and lane 4: 100 bp ladder.

These results indicate that the mutation is likely to lead to skipping of exon during splicing event (Figure 2). To confirm the results from RNA assay, sequencing of specific bands form RT_PCR for ARMC9_pCAS_Wt and ARMC9_pCAS_Mt was done by Sanger sequencing, which revealed deletion of 99 base pairs of exon 9 of *ARMC9* gene (Supplementary Figure 1). This result confirms that synonymous mutation c.879G>A leads to abolition of splice site and leads to skipping of exon 9 of *ARMC9* which may result in in-frame deletion of 33 amino acids (exon 9) form *ARMC9* in patients.

## DISCUSSION

We elucidated the molecular basis in three siblings who were diagnosed with mental retardation, ptosis and polydactyly phenotype, using homozygosity mapping and whole exome sequencing. Whole exome sequencing analysis revealed a homozygous synonymous splice site variant in *ARMC9* gene (c.879G>A) in these patients. This variant leads to abolition of splice site, which leads to skipping of exon 9 during splicing event.

Intellectual disability in combination with other clinical features represents a heterogeneous group of genetic disorders with large number of causal genes^7^. Combination of homozygosity mapping and exome sequencing has allowed us to examine small family with few affected individuals, particularly in the case of recessive mode of inheritance and consanguinity, to efficiently limit the number of candidate genes to be studied and establish new disease-gene association.

*ARMC9* was first reported in cDNA analysis of human fetal whole brain, which revealed expression of several new long coding transcripts. One of them was *ARMC9*, which was reported as KIA1868^8^. ARMC9 is also known as LisH domain containing protein ARMC9, NS21^9^, Melanoma/melanocyte specific tumor antigen KU-MEL-1^9^ and KIAA1868. ARMC9 is a lesser known protein of 817 amino acids, containing N-terminal LisH (Lissencephaly type 1like homology) domain and C-terminal ARM (Armadillo repeats) motifs. Proteins containing LisH domains are known to be involved in microtubule dynamics and cell division^10^. Armadillo (ARM) repeats are characterized as repeating, 42 amino acid motifs, composed of three α helices. Although ARM repeat containing proteins do not share sequence similarity, they share a similar structure^11^. Multiple ARM repeats fold together to form a super-helix structure, which acts as platform for interaction with protein partners. ARM containing proteins are conserved through eukaryotes and are involved in diverse cellular functions such as signal transduction, cell migration and proliferation and cytoskeletal regulation^12, 13^.

We proceeded further to investigate ARMC9 function based on homology. We could not detect clear homolog but BLAST (https://blast.ncbi.nlm.nih.gov/Blast.cgi?PROGRAM) and InterProScan^14^ protein domain homology searches showed that there are two predicted ARM domains (154-341 and 375-584) at the C-terminus, in addition to N-Terminal LiSH domain. The tandem ARM repeat domains of ARMC9 may fold together as a series of tandem helices forming a super-helix that creates a surface or groove for protein interaction similar to that of the Beta catenin (CTNNB1) ARM repeat structure. Yeast two hybrid assay has shown that ARMC9 interacts with Siah E3 ubiquitin protein ligase 1 (SIAH1) and CKLF like MARVEL transmembrane domain containing 5 (CMTM5)^15^, which indicates that ARMC9 may be involved in ubiquitination pathway like ARMC8^16^.

To study effect of mutation in splicing, the best method is analysis of patient RNA for respective gene, but *ARMC9* has little or no expression in adult tissue.^9^ Thus we proceeded with alternate method for assay of splicing defect using PCAS2 miningene system. Synonymous splicing defect mutation c.879G>A, which leads to skipping of exon 9 in *ARMC9* gene will lead to in-frame deletion of few ARM repeats in ARM domain (deletion of 260-293 aa), that is likely to influence protein binding capabilities of ARMC9. It is known that armadillo repeat containing proteins can have more than one function in cells, potentially interacting with different protein partners^17^.

Several studies on another ARM contacting protein Beta catenin (CTNNB1) indicate that mutations in *CTNNB1* can lead to wide range of neurodevelopmental disorders (ID with syndromic features^18, 19^. Studies on mouse mutant that carries mutation in armadillo repeat of Beta catenin (*CTNNB1*) demonstrate reduced affinity for membrane associated cadherins^20^. Similarly mutation in *APC2*, an ARM containing protein involved in neural development, is associated with Sotos syndrome 3 (MIM: 61719) involving ID, hyperactive behaviour with macrocephaly ^21^. Thus ARMC9 also is likely to be involved in neurodevelopmental process and its perturbation may lead to syndromic ID.

In summary, here we report on a synonymous splice site variant in *ARMC9* as a cause of mental retardation, ptosis and polydactyly phenotype. The variant affects conserved ARM repeats at the C terminus and is likely to disrupt its interaction with other protein partners. *ARMC9* joins an important group of highly conserved ARM repeat containing proteins associated with intellectual disability which includes Beta catenin (*CTNNB1*), *APC2*. The exact nature and essential role of *ARMC9* remains to be fully characterized, however these results further expand our understanding of molecular genetic basis of intellectual disability and facilitate better counselling of patients.

## ACKNOWLEDGEMENTS

We are grateful to the family for their support during this study. We acknowledge MedGenome (Cochin, Kerala, India) for performing exome sequencing. We also acknowledge the funding support from Department of Biotechnology, Government of India, Grant Number: BT/PR3193/MED/12/521/2011 and Indian Council of Medical Research (BMS 54/2/2013).

Supplementary information accompanies this paper on European Journal of Human Genetics website.

### Web resources

1000 Genomes – http://www.1000genomes.org/

Exome Variant Server – http://evs.gs.washington.edu/EVS/

ExAC – http://exac.broadinstitue.org/

dbSNP- http://www.ncbi.nlm.nih.gov/SNP/

ClinVar – http://www.ncbi.nlm.nih.gov/clinvar/

OMIM – http://www.omim.org/

HGMD – http://www.biobase-international.com/products/hgmd

Integrative Genomics Viewer (IGV) - http://www.broadinstitute.org/igv/

SIFT – http://sift.jcvi.org/

Polyphen2 – http://genetics.bwh.harvard.edu/pph2/

HANSA – http://www.cdfd.org.in/HANSA/

